# Fitness seascapes are necessary for realistic modeling of the evolutionary response to drug therapy

**DOI:** 10.1101/2022.06.10.495696

**Authors:** Eshan S. King, Anna E. Stacy, Davis T. Weaver, Jeff Maltas, Rowan Barker-Clarke, Emily Dolson, Jacob G. Scott

## Abstract

Pharmacokinetic (PK) and pharmacodynamic (PD) modeling of host-pathogen interactions has enhanced our understanding of drug resistance. However, how combinations of drug resistance mutations impact dose-response curves remains underappreciated in PK-PD studies. The fitness seascape model addresses this by extending the fitness landscape model to map genotypes to dose-response functions, enabling the study of evolution under fluctuating drug concentrations. Here, we present an empirical fitness seascape in *E. coli* harboring all combinations of four drug resistance mutations. Incorporating these data into PK-PD simulations of antibiotic treatment, we find that higher mutation supply increases the probability of resistance, and early adherence to the drug regimen is critical. In vitro studies further support the finding that the second dose in a drug regimen is important for preventing resistance. This work represents the first application of an empirical fitness seascape in computational PK-PD studies, revealing novel insights into drug resistance.

---

Antimicrobial resistance (AMR) is a persistent challenge that contributes to substantial global mortality and economic burden, with disproportionate impacts on those in low-resource settings (*1*). Understanding and predicting the emergence of drug resistance within and among human hosts is an intuitive path towards developing novel effective treatment strategies. The emergence of resistance is studied through an evolutionary lens, whereby novel genotypes of a disease agent emerge through a random process and are selected for due to their increased fitness. Recently, computational studies of the evolution of resistance have been used to optimise dosing schedules *in silico*, *in vitro*, and in clinical trials (*2–9*).

To understand, simulate, and predict evolution, researchers often use the fitness landscape model, which maps genotype to fitness (*10–16*). An assumption of the fitness landscape model is that, for a given landscape, the selection pressure is constant (*12*). As such, canonical fitness landscapes cannot model evolution under continuously varying selection pressure, such as the rising and falling concentration of drug in a patient over time. This limitation of singular fitness landscapes precludes the modeling of evolutionary trade-offs, or fitness costs, where a mutant “trades” a lower growth rate in the absence of a drug in exchange for a higher growth rate at higher concentrations of the drug (*17–19*).

Such trade-offs across environments are common in evolutionary medicine; drug resistance mechanisms often impose metabolic burdens or impair vital functions of the organism. When the drug selective pressure is removed, the sensitive genotype may have a higher relative fitness and the resistant genotype may go extinct (*20, 21*). Moreover, disease agents in a patient will never experience a constant environment – the drug concentration will vary according to the pharmacokinetic profile, dosing schedule, and spatial distribution. Fitness seascapes extend the fitness landscape model by mapping both genotype and environment (i.e. drug concentration) to fitness (*22–28*). Here, we model fitness seascapes as collections of genotype-specific dose-response curves (**Fig 1A**), similar to our previous work (*28*). This approach permits the modeling of fitness trade-offs across a range of drug concentrations, allowing us to model the evolution of resistance with realistic pharmacological considerations.

**Figure 1:**
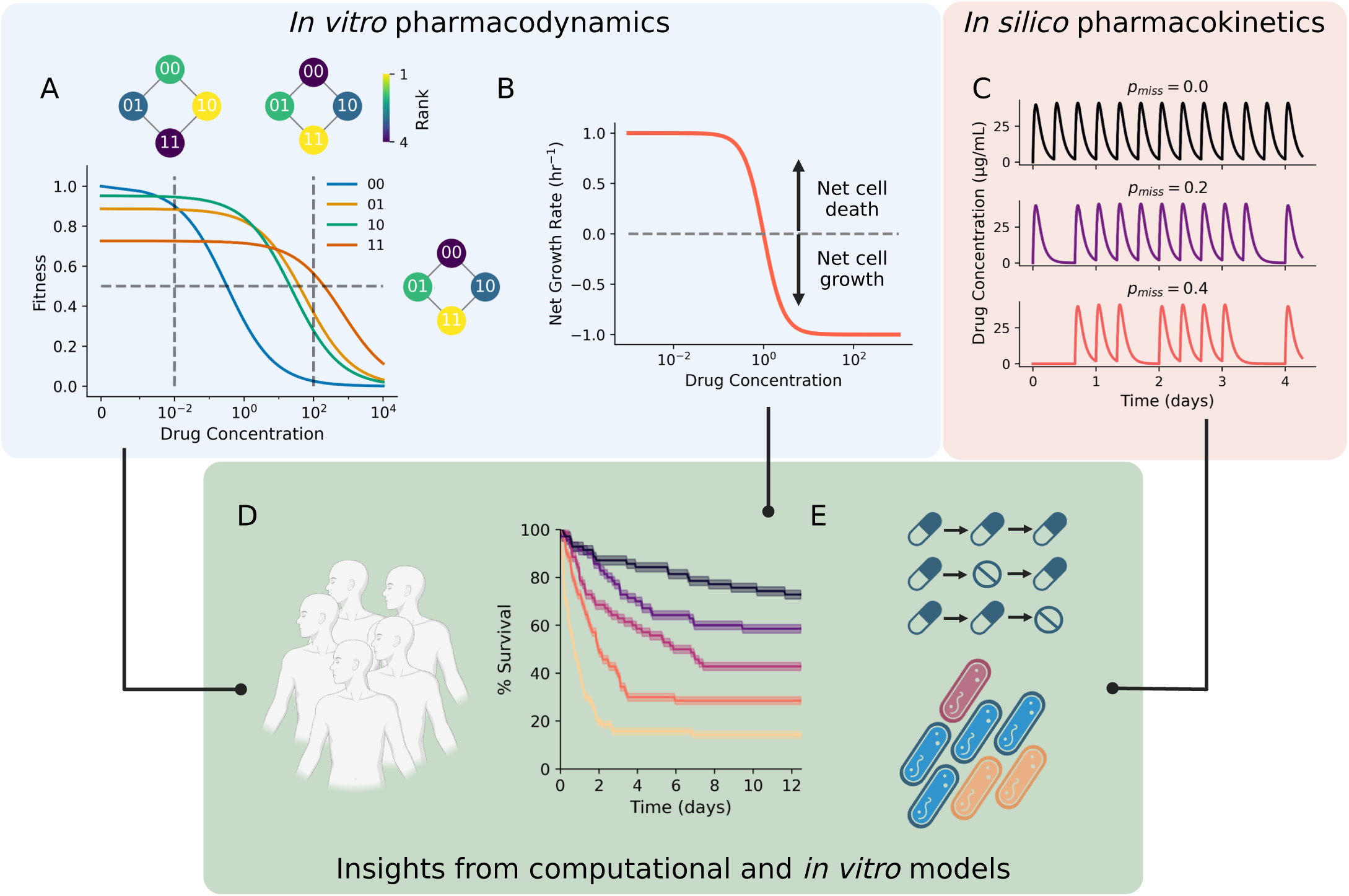
Fitness seascapes intrinsically model evolutionary tradeoffs. (**A**) Fitness seascape parameterized with random synthetic data. The genotype with the highest IC_50_ (11) also exhibits the lowest drug-free growth rate, suggesting a tradeoff, or cost, to resistance. Rank-order fitness landscapes based on growth rate are shown inset, with the vertical dotted line at the corresponding drug concentrations. A fitness landscape based off of IC_50_ alone is also shown with a corresponding horizontal line. (**B**) Time-kill experiments are used to parameterize bacterial dose-response curves that include net cell death (net growth rate less than zero). (**C**) Patient dosing regimens are modeled with pharmacokinetic curves, which can be used to study patient nonadherence. The probability of a simulated patient missing a scheduled drug dose is given by *p*_*miss*_. (**D**) In this work, we simulate the pharmacokinetic characteristics of a wide variety of patients and study how infectious bacterial populations respond. Simulations are parameterized by integrated *in vitro* fitness seascapes and incorporate *in silico* pharmacokinetic curves. (**E**) We use *in silico* and *in vitro* models to evaluate the impact of drug dosing nonadherence on the probability of drug resistance.

In this work, we describe a novel empirical fitness seascape in a clinically-relevant *E. coli* drug resistance model. We show that drug resistance in this model is associated with a substantial fitness tradeoff, with a higher IC_50_ associated with a lower drug-free growth rate. We use this seascape and empirical dose-dependent death rates estimated from a time-kill assay to parameterize a computational model to explore IV and oral dosing regimens. In the IV antibiotics setting, we simulated patients treated for *E. coli* bacteremia, which carries a high rate of treatment failure with mortality estimates between 12% and 33% (*29–31*). In the outpatient setting, we simulated patients treated with oral antibiotics for less severe infections such as a urinary tract infection or pneumonia. We find that pathogen parameters such as mutation rate and population size impact the probability of treatment success in IV antibiotics regimens. We explore patient drug regimen nonadherence with oral dosing regimens and find that early drug regimen adherence is important for treatment success. We further validate this result by modeling treatment nonadherence *in vitro*, finding that missing the second dose results in treatment failure. Our work represents a novel application of an *in vitro* fitness seascape for studying clinical drug dosing regimens, and serves as a bridge between evolutionary medicine and clinically-relevant PK-PD models.

## Fitness seascapes as a model for evolutionary tradeoffs

Using synthetic data, we first explored how evolutionary tradeoffs may be examined with fitness seascapes. Here, we define “tradeoff” as a case where a genotype with a higher IC_50_ grows slower in the absence of drug than a genotype with a lower IC_50_. We also use “cost of resistance” interchangeably with tradeoff. **Fig 1A** shows a synthetic fitness seascape that illustrates fitness tradeoffs: genotype 00 has the highest drug-free growth rate and exhibits the lowest IC_50_, while genotype 11 has the lowest drug-free growth rate and the highest IC_50_. This represents the case where drug resistance mechanisms impose some metabolic burden on the organism, such as the upregulation of a protein or more energetically costly synthesis. Notably, tradeoffs induce changes to the fitness landscape as the drug concentration changes, as shown in the inset rank-order fitness landscapes based on growth rate. In contrast, the inset fitness landscape based on IC_50_ alone cannot intrinsically model tradeoffs in this way.

## *In-vitro* fitness seascape reveals evolutionary tradeoffs in *E. coli*

Genetically-engineered *E. coli* cell lines were provided by the Weinreich lab at Brown University (*10*). Details of the model system are provided in methods. Briefly, a combinatorially-complete set of clinically-relevant drug resistance point mutations was introduced into the *bla* gene encoding the ampicillin-resistance (Amp_R_) β-lactamase protein. Sixteen genotypes were generated by engineering all possible combinations of four point mutations. We denote the presence or absence of a point mutation with a 0 (absence) or 1 (presence) in a binary string of length four, with each position in the string corresponding to a unique point mutation. 0000 denotes the wild-type *bla* gene; 0001 represents G238S, 0010 represents M182T, 0100 represents E104K, and 1000 represents A42G. Each of these point mutations is in the penicillin binding pocket of β-lactamase and serves to increase the rate of cefotaxime hydrolysis (*32*).

We quantified dose-response curves for each genotype by estimating growth rate as a function of drug concentration using an optical density (OD) assay (details in methods). The degree of drug resistance and drug-free growth rate vary widely across the 16 genotypes (**Fig 2**). To understand the potential fitness costs to resistance-conferring mutations, we investigated the relationship between the number of mutations and the drug-free growth rate, finding that the two were inversely related (**Fig 2B**; *p* = 0.003, *r*^2^ = 0.48). We also found a positive correlation between the number of mutations and IC_50_ (*p* = 0.004, *r*^2^ = 0.46) and an inverse relationship between drug-free growth rate and IC_50_ (*p* = 0.011, *r*^2^ = 0.38) **Fig 2C** and **Fig 2D**).

**Figure 2:**
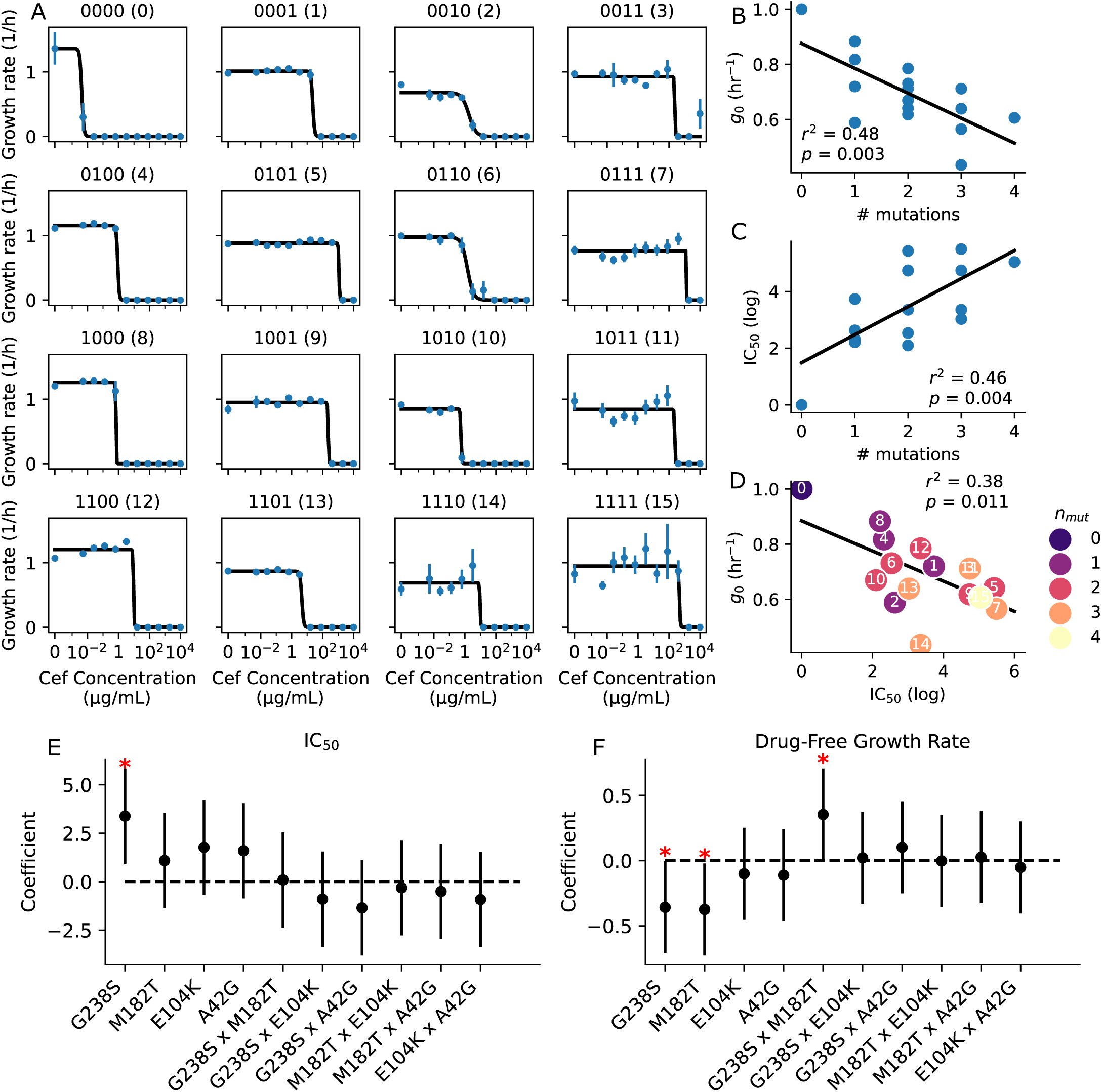
*In vitro* pharmacodynamics reveals fitness costs to resistance. (**A**) Cefotaxime dose-response curves for all 16 genotypes. Growth rate estimates with standard deviation are shown in blue (*N* = 8 replicates per condition). Pharmacodynamic curve fit is shown in black. (**B**) Drug-free growth rate (*g*_0_) versus the number of mutations for all 16 genotypes. Linear regression is shown in black (*r*^2^ = 0.48, *p* = 0.003). (**C**) IC_50_ versus number of mutations. Linear regression is shown in black (*r*^2^ = 0.46, *p* = 0.004). (**D**) Drug-free growth rate versus IC_50_. Color of each data point indicates the number of mutations, while the number inset corresponds to the genotype. Linear regression is shown in black (*r*^2^ = 0.38, *p* = 0.011). (**E**) Multivariate regression assesing the impact of all four mutations and pairwise combinations on IC_50_. Red asterix indicates significant factor. Black bars indicate 95%-ile interval. (**F**) Multivariate regression assesing the impact of all four mutations and pairwise combinations on IC_50_. Red asterix indicates significant factor. Black bars indicate 95%-ile interval.

We next investigated whether particular resistance mutations were more costly than others, and, similarly, if different resistance mutations conferred more resistance than others. To do this, we performed a multivariate linear regression with each resistance mutation as an independent predictor and either drug-free growth rate or IC_50_ as the outcome. We also included all pairwise interaction terms in the analysis. Multivariate analysis revealed that G238S was the only mutation independently associated with IC_50_ (**Fig 2E**). Interestingly, we found that both G238S and M182T were independently negatively associated with drug-free growth rate while the interaction term G238SxM182T was positively associated with drug-free growth rate (**Fig 2F**). These results suggest a reciprocal sign epistasis, where G238S and M182T are both deleterious on their own but advantageous to drug-free growth rate when combined.

## Estimating dose-dependent death rates with a time-kill assay

We sought to apply our *E. coli* fitness seascape to computational studies of clinical drug dosing by modeling populations of bacterial cells treated with an antibiotic. In order to fully parametrize an agent-based algorithm, a robust model for cell death was necessary. However, experimental quantification of cell death rates was challenging. We found that we could not observe dynamics above the minimum inhibitory concentration (MIC) with our optical density data – in the above-MIC range, we simply saw no cell growth. Previously published time-kill methods have several drawbacks, such as requiring hundreds of cell culture plates or suffering from residual drug carrying over into the cell count assay. To improve upon the current status-quo and better parameterize drug dose-dependent death rates, an accurate, facile, and low-footprint time-kill assay is needed.

To address these shortcomings, we developed a fluorescence-based assay that utilizes resazurin as a cell-viability indicator (details in methods). Resazurin is a dye that is blue and weakly fluorescent and is rapidly reduced to the pink and brightly fluorescent resofurin by viable cells. Briefly, a cell-count to fluorescence calibration curve was generated and then used to estimate cell count over time for cell cultures with different concentrations of cefotaxime. Ten time points were taken for each condition over the course of ~5 hours, with three technical replicates per condition. Summary cell count over time data is shown in **Fig 3A**. We then estimated the maximum net change in cells, referred to as the “net growth rate”, for each drug concentration and fit these data to a pharmacodynamic curve (**Fig 3B**). Finally, we combined this pharmacodynamic relationship with the drug-free growth rate and IC_50_ estimated in **Fig 2A** to obtain dose response curves with positive and negative net growth rates.

**Figure 3:**
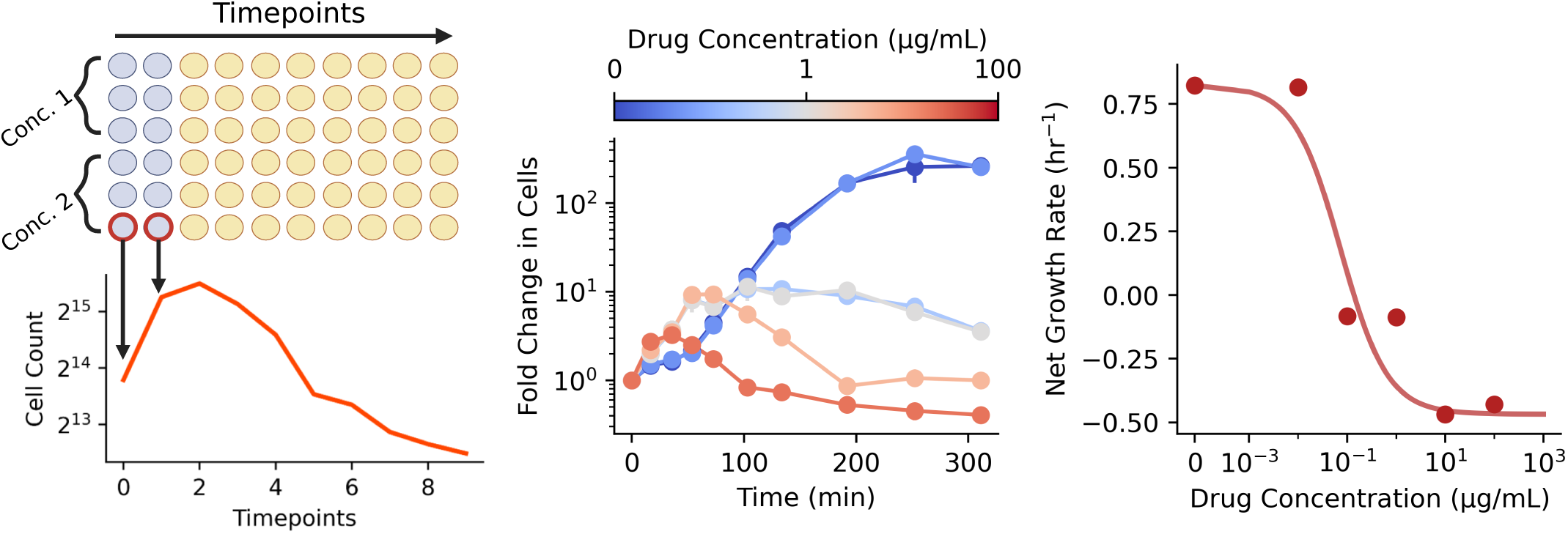
A novel time-kill assay for estimating dose-dependent death rates. **(A)** Illustration of the fluorescence-based time-kill assay used to parameterize bacterial death rates. Circles represent wells with bacterial culture in a 96-well plate; blue circles represent wells with alamarBlue (AB) added. Cell count over time for a single replicate is shown below in orange. **(B)** Cell count over time estimated from a time-kill assay for different concentrations of cefotaxime. Standard deviation bars may be smaller than the plotted data point. *N* = 3 replicates for each timepoint. **(C)** Growth rate versus drug concentration estimated from data in **B**. Solid line shows pharmacodynamic curve fit. Standard deviation bars are smaller than the plotted data point.

## Pathogen mutation rate and population size impact the probability of treatment success

To understand what factors could contribute to treatment failure in *E. coli* infections, we applied our fitness seascape and time-kill data to computational models. To do this, we simulated patients undergoing a 10-day therapy regime with cefotaxime with an agent-based model: Fast Evolution on Arbitrary Seascapes (FEArS). FEArS models the number of dying, dividing, and mutating cells with a Poisson process (**Fig. S1**). Mutations are modeled with an agent-based algorithm whereby mutating cells are randomly allocated to “adjacent” genotypes (genotypes that differ by Hamming distance 1). We parameterized FEArS with the fitness seascape and time-kill data described above to simulate infections with our *E. coli* model system. Agent growth rate in the sub-MIC regime (i.e. positive net growth rate regime) was parametrized with growth rate data from the fitness seascape, while growth rate in the above-MIC regime was parameterized with time-kill data. We modeled patient serum pharmacokinetic profiles with a one-compartment model:

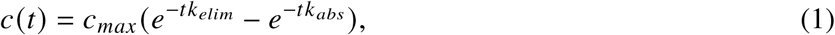

where *c*(*t*) is the serum concentration over time, *c*_*max*_ is the maximum achieved concentration, *k*_*elim*_ is the drug clearance rate, and *k*_*abs*_ is the drug absorption rate. *N* doses were modeled as a Dirac comb with the period *T* equivalent to the dosing frequency:

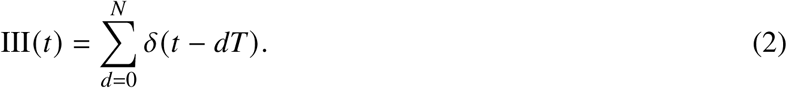

The final patient pharmacokinetic profile is then

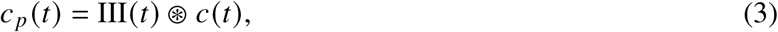

where ⊛ is the convolution operator.

Patient pharmacokinetics differ most notably in the maximum serum concentration *c*_*max*_ and the drug half life *t*_1/2_, which is governed by *k*_*elim*_ in **Eq 1** (*33*). Infections can also differ in their mutation rate *r*_*m*_ (*34,35*) and population size, here modeled as a carrying capacity *K*. To examine the interaction of all of these parameters, we used Latin Hypercube Sampling (LHS) to generate 300 sets of each of these parameters (*36*). LHS allows for uniform random sampling of a high-dimensional grid, ensuring that a wide and representative range of parameters and parameter combinations are tested. For each set of four parameters, we ran 100 simulations and computed the rate of treatment success (0 surviving cells by the end of the simulation). Parameter ranges tested are shown in **Table 1**. Interestingly, *c*_*max*_ and *t*_1/2_ had no discernible effect on treatment efficacy, while *K* and *r*_*m*_ both strongly impacted treatment outcome (**Fig 4A**). Multiple linear regression analysis revealed that there was a strong interaction between *K* and *r*_*m*_ (*β* = 1.46); a summary of the linear regression coefficients is shown in **Table S3**. We visualized this interaction by plotting the pairwise joint distributions between the parameters (**Fig 4B**), finding that increasing *K* and *r*_*m*_ jointly decreased the probability of success.

**Figure 4:**
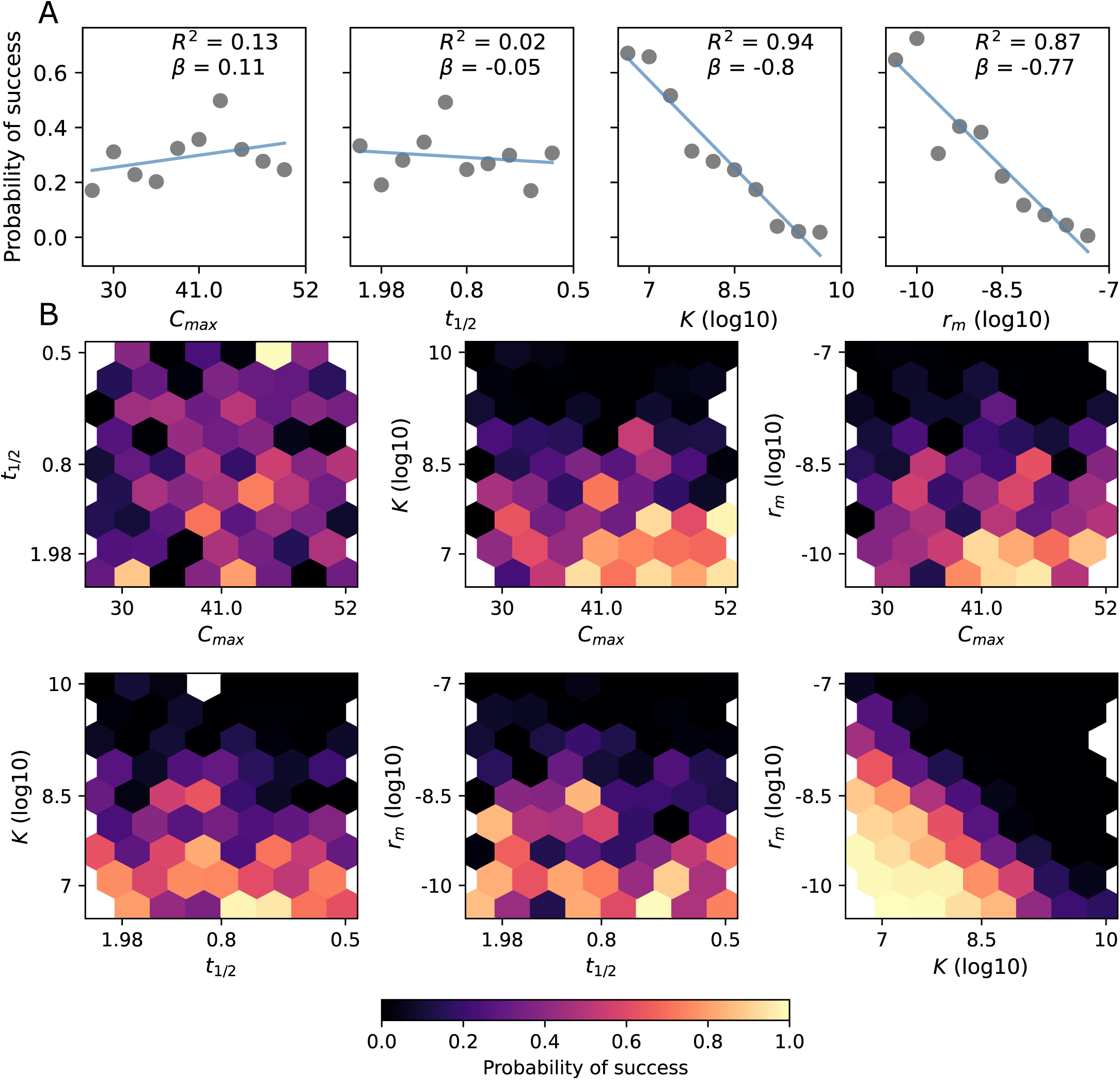
Impact of pharmacokinetics, carrying capacity, and mutation rate on treatment success. **(A)** Marginal distributions of probability of treatment success versus *t*_1/2_, *C*_*max*_, *r*_*m*_, and *K* for *N* = 100 simulated patients per condition. Linear regression for each parameter is shown in blue. The coefficient of determination *R*^2^ and coefficient of the linear regression *β* are shown inset. **(B)** Joint distributions of probability of treatment success. Treatment success was defined as 0 cells surviving by the end of the simulation. Dose frequency was one dose per 8 hours, duration of treatment was 10 days, and the simulations were allowed to reach equilibrium for 3 days before beginning treatment. Latin hypercube sampling was used to generate 300 sets of parameters.

**Table 1:** Parameter ranges used for simulations with literature sources.

| Parameter | Description | Value Range | Citation |
| --- | --- | --- | --- |
| $c_{max}$ | maximum serum concentration | 30-52 $\mu\text{g/mL}$ | (33) |
| $K$ | carrying capacity | $10^7 - 10^{10}$ cells | n/a |
| $t_{1/2}$ | drug half-life | 0.5-2 hr | (33) |
| $r_m$ | mutation rate | $10^{-10} - 10^{-7}$ bp$^{-1}$ | (34, 35) |

We further examined the interaction between *K* and *r*_*m*_ by plotting the rate of treatment success against average mutation supply, *K* ∗ *r*_*m*_, revealing a steep drop off in treatment success at ~1 mutant per generation (**Fig S2**). Notably, prior work has found that high-mutation rate phenotypes are common among pathogenic *E. coli* (*37*) and high mutation rate is strongly correlated with drug resistance in the clinic (*38*). Analyzing the evolutionary trajectory of the simulations revealed that almost all simulations in which there was treatment failure acquired the G238S mutation (allele 0), allowing for evolutionary escape (**Fig S3**). G238S is the mutation that confers the greatest change in IC_50_, on average, and has been found in many clinical samples. A high mutation supply produced a small population of genotype 1 (0001) and 2 (0010) in equilibrium with genotype 0 prior to the onset of therapy, despite there being a fitness cost.

## Patient nonadherence contributes to treatment failure

Patient nonadherence is thought to contribute to treatment failure and antibiotic resistance in the ambulatory setting (*39–41*); studies have found that 30-45% of patients are nonadherent with their antibiotic regimen, with a large proportion of the nonadherence due to forgetfulness (*39, 41*). To investigate this, we modeled varying drug elimination half life *t*_1/2_, time to maximum drug concentration *t*_*max*_, and drug regimen nonadherence. We calculated the appropriate *k*_*elim*_ and *k*_*abs*_ from *t*_*max*_ and *t*_1/2_ and again applied **Eq 1** to model patient serum drug concentration. Patient nonadherence was modeled as a random process with each patient having a constant probability *p_forget_*of missing a scheduled dose. We modeled patients undergoing a 7-day treatment regimen using FEArS and measured the probability of treatment success for different combinations of the three parameters (*N* = 100 simulations per condition). Interestingly, there was a significant additive interaction between *t*_1/2_ and *p_forget_* (**Fig 5A**), with increasing *p_forget_*and decreasing *t*_1/2_ jointly decreasing the rate of treatment success. In contrast, there was no clear relationship between *t*_*max*_ and treatment success and no interaction between *p_forget_* and *t*_*max*_. In order to better understand what drives treatment failure in drug regimen nonadherence, we fixed *t*_1/2_ and *t*_*max*_ (6 and 1 hr, respectively) and simulated patients with varying rates of *p_forget_*(*N* = 1000 patients per condition). Representative drug concentration curves are shown in **Fig 5B**. We found that different rates of *p_forget_* resulted in dramatically different time-to-event (TTE) curves, with higher rates of nonadherence leading to both lower rates of success and longer times to treatment success (**Fig 5C**).

**Figure 5:**
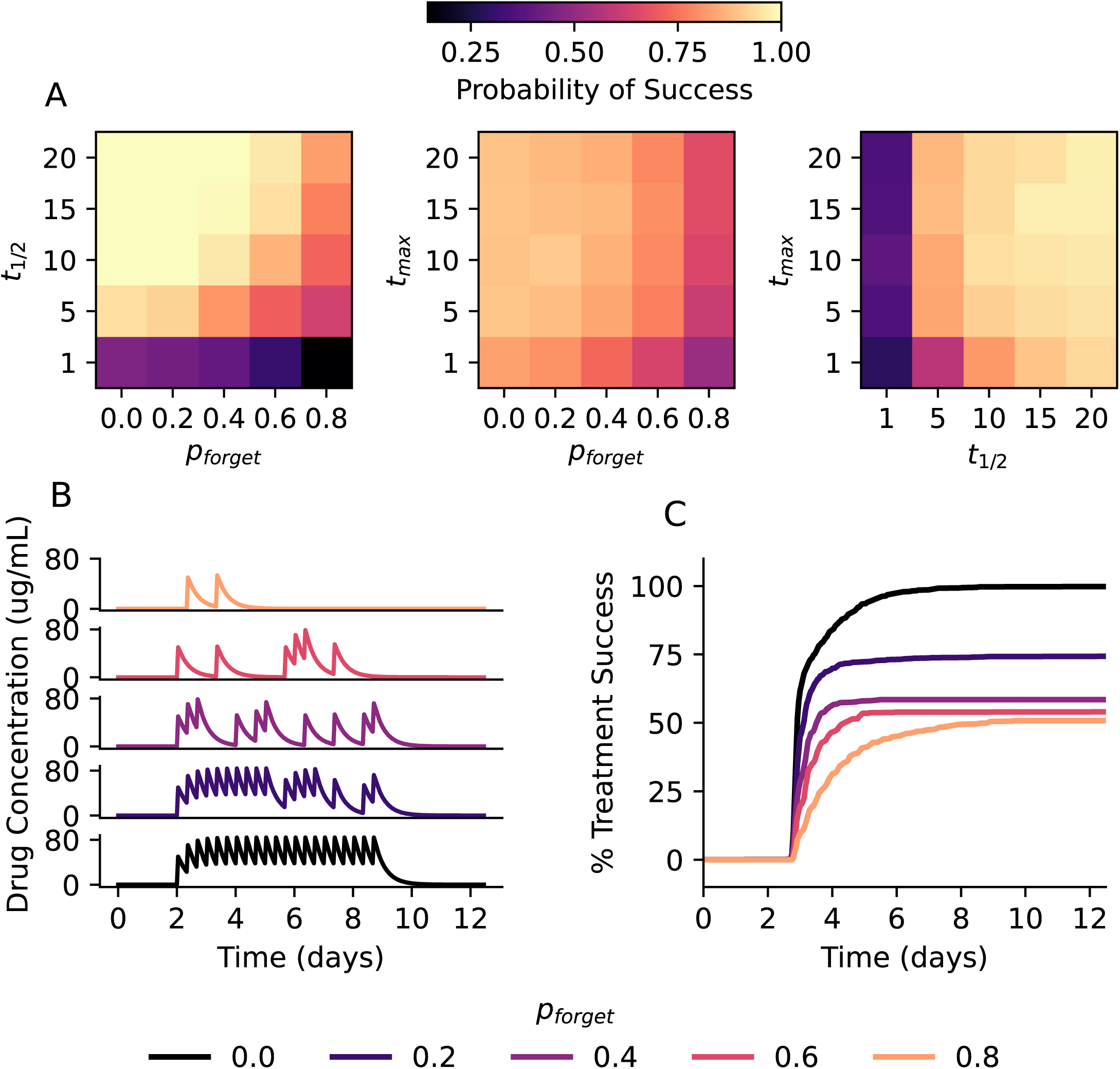
Patient nonadherence and drug half-life impact treatment success. **(A)** Joint distributions of probability of treatment success for *p_forget_*, *t*_1/2_, and *t*_*max*_. *N* = 100 simulations per condition. **(B)** Example simulated patient pharmacokinetic curves for different values of *p_forget_*. **(C)** Time-to-event curves of treatment success for different rates of *p_forget_*. *N* = 1000 simulations per *p_forget_*. 95% confidence intervals are smaller than the widths of the plotted lines.

We hypothesized that there may be patterns in patient nonadherence that impact treatment success. We calculated the risk ratio of each individual drug dose and found that, for moderate rates of nonadherence (*p_forget_* <= 0.6), early drug doses were particularly important for treatment success (**Fig 6A**). In this case, a higher risk ratio indicates that a particular dose was more important for treatment success. Based on this finding, we calculated the probability of treatment success partitioned by time between the first two doses, either equal to 8 hours or greater than 8 hours between the first two doses (**Fig 6B**). We found that regimens where the second dose was administered exactly 8 hours after the first had a higher rate of success compared to regimens where the second dose came greater than 8 hours after the first, and this difference increased as the nonadherence rate decreased. This suggests that at low rates of nonadherence, appropriately timing the first two doses is critical for treatment success. We also tested if the total number of doses correctly administered corresponded to treatment success for a given *p_forget_*, finding that the average difference in total doses between successful and unsuccessful regimens was less than 1 (**Fig S4**). This observation suggests that the total number of doses is not a meaningful independent predictor of treatment success.

**Figure 6:**
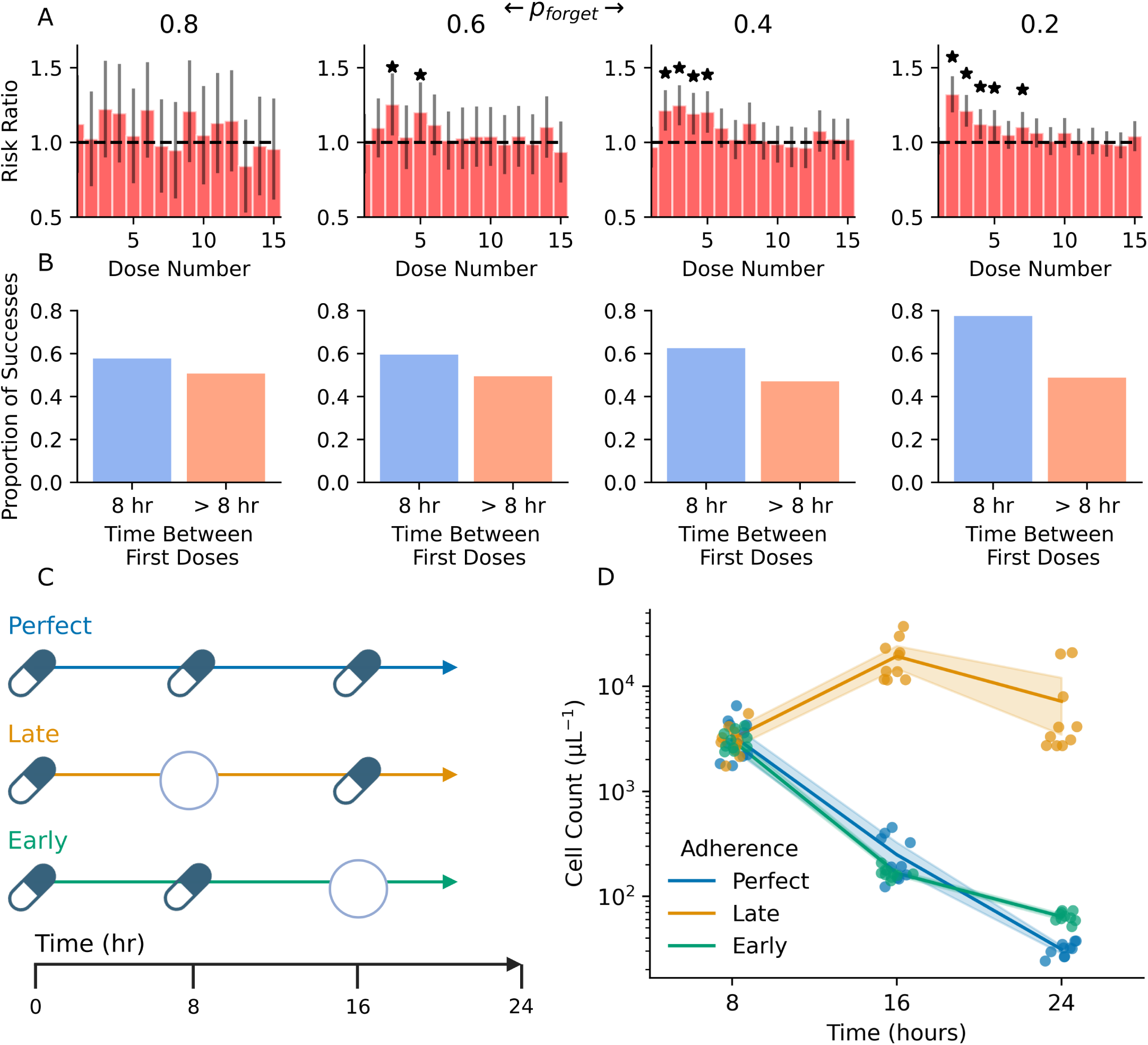
Early drug regimen adherence contributes to treatment success. Each column corresponds to a different *p_forget_*, labeled at the top. **(A)** Risk ratio for each dose based on the probability of treatment success. A higher risk ratio indicates a stronger association with treatment success. Error bars indicate 95% confidence intervals. Stars indicate where the 95% CI excludes 1. **(B)** Proportion of treatment success partitioned by regimens with 8 hours between the first two doses (blue column) and greater than 8 hours between the first two doses (orange column). *N* = 1000 simulations per *p_forget_*. **(C)** Illustration of *in vitro* drug dosing experiment. Perfect refers to all three correctly administered drug doses; late refers to a missed second dose and correctly administered third dose (i.e. late adherence); early refers to a correctly administered second dose and missed third dose (i.e. early adherence). **(D)** Cell count over time estimated with AB for three conditions in **(C)**. Shaded region shows 95% CI for each timepoint. *N* = 8 replicates per condition.

## *In vitro* validation supports importance of early adherence

In order to further explore the computational finding that early nonadherence impacts treatment success, we modeled nonadherence *in vitro* and studied the impact of different dosing regimens. Using co-cultures of genotypes 0, 1, and 2 to generate standing genetic heterogeneity, we dosed cultures in 96-well plates with three different cefotaxime regimens (**Fig 6C**): perfect, representing 100% adherence (3 successive doses with 8 hour intervals); late, representing late adherence (i.e. missing the second dose); and early, representing early adherence (i.e. missing the third dose). Cultures were passaged every 8 hours and the drug dose was adjusted during passaging according to the dosing regimen. Note that the cells were not re-suspended in fresh media during passaging, so there was some drug carry-over; this results in a ∼1 μg/mL difference in drug concentration between the perfect adherence regimen and the nonadherence regimens, and likely does not affect the qualitative interpretation of the experiment. After each 8 hour interval and before passaging, plates were scanned with AB to estimate cell count over time. In the perfect adherence condition, the cell count dropped continuously after each successive dose, demonstrating robust treatment response (**Fig 6D**). In the early adherence condition, the cell count mirrored that of the perfect adherence condition despite missing the third dose at 16 hours, suggesting that the two early doses strongly reduced the viable cell count. In contrast, in the late adherence condition, there was continued cell growth after the third scheduled dose – after missing the second dose, the population rebounded to ~10^4^ cells/μL. Following the third administered dose at 16 hours, the population count did not fall appreciably, demonstrating drug resistance. These results support the finding that two successive doses at the beginning of the regimen are vital to prevent drug resistance and successfully inhibit population growth.

## Discussion

Here, we have described a novel, *in vitro* fitness seascape of a clinically-relevant drug resistance model and used it to generate insights into clinical drug dosing regimens. Importantly, by quantifying the growth rate of genotypes representing a combinatorially-complete set of drug resistance mutations, we have found that resistance-conferring mutations in the TEM β-lactamase gene impose substantial fitness costs in *E. coli*. This finding aligns closely with a meta-analysis of fitness costs to drug resistance in bacteria, where the magnitude of the fitness cost was found to be positively correlated with the minimum inhibitory concentration (MIC), which is an indicator of drug resistance (*42*). Allele-level analysis revealed that G238S and M182T are both negatively correlated with drug-free growth rate, but the interaction term G238SxM182T was positively correlated, suggesting reciprocal-sign epistasis. This interaction may relate to the biochemical function of the M182T mutation, which stabilizes other drug-resistance mutations in TEM including G238S. This finding may also help explain why G238S and M182T are often found together in clinical TEM isolates (*43*). We also found that G238S is significantly associated with IC_50_ – previous biochemical and computational studies of the TEM β-lactamase have shown that the G238S mutation is associated with the highest increase in cefotaxime hydrolysis and is likely a major driver of clinically-relevant drug resistance (*44*). Together, these genotype- and allele-level findings provide novel insights into the cost of drug resistance in bacteria.

We used this seascape model to explore intravenous and oral drug dosing regimens with computational agent-based modeling. We found that parameters that contribute to mutation supply – mutation rate and population size – most strongly influence the probability of treatment success in our model of *E. coli* bacteremia. In the simulated outpatient setting with oral antibiotic dosing, we found that drug regimen nonadherence is associated with treatment failure, but early drug regimen adherence contributes to treatment success. We further validated this finding *in vitro* by studying the impact of the time between drug doses, finding that missing the second dose in a series (i.e. “late” adherence) strongly correlates with evolutionary escape.

Mathematical and computational modeling of within-host evolutionary dynamics has a long and rich history, especially in applications to long-term infections such as HIV and tuberculosis (*45–48*). Here, we have expanded on this prior work in several key ways. First, we parameterized our simulations with an empirical fitness seascape, enabling us to study evolutionary dynamics with a clinically relevant drug resistance model. By quantifying growth rates, and thus inherently capturing fitness tradeoffs, fitness seascapes provide a more realistic model for selection under varying drug concentrations. In addition, we developed a novel time-kill assay to estimate dose-dependent bacterial death rates, allowing us to more accurately simulate cell death in our agent-based models. Combining empirical genotype-specific growth rates and drug dose-dependent death rates reduces the number of free biological parameters in our model, increasing the external validity of our findings. Furthermore, we used a hybrid agent-based algorithm (FEArs) for stochastic simulations. This choice enabled efficient computational experiments, allowing for a large number of parameter combinations and replicates while maintaining important biological constraints, even for very small population sizes. This innovation allows us to examine population extinction and evolutionary rescue from extinction, which is important for modeling the emergence of drug resistance in the clinic. In addition to the methodological advances, our work provides insights into the somewhat underappreciated evolutionary dynamics of short-term infections in the clinic. Our findings suggest that the emergence of *de novo* drug resistance mutations during the course of therapy may contribute to treatment failure in both the inpatient and outpatient settings with timescales of days to weeks.

A natural extension of this work is in modeling antibiotic cycling to avoid resistance in bacteria. Much interest has been given to optimizing antibiotic cycling in both computational studies with fitness landscapes (*9, 11, 49–52*), *in vitro* studies (*53*), and clinical trials (*54*). In these models, different antibiotics are associated with different fitness landscapes. Each landscape is, in essence, a tool that can be used to push an evolving population in a desirable direction. By conceptualizing these landscapes as seascapes, we can simultaneously increase the accuracy of our models and the number of distinct fitness landscape topographies available to use for the purposes of steering evolution (*51, 55*). Each individual drug may produce several different changes in genotype fitness rank as a function of antibiotic concentration.

Optimizing drug dosing regimens is another natural application of our work. By varying the timing and dose of an antibiotic therapy, we may discover parameter ranges that reliably avoid resistance *in silico*. Then, using empirical knowledge of patient preferences (*56, 57*), we may deliver dosing regimens that are both amenable to patient adherence and successful elimination of the infection.

There are several important limitations to this work. The 4-allele fitness seascape quantified here represents only a small set of possible resistance mutations. For instance, off-target mutations in genes regulating drug efflux pumps or cell metabolism may also contribute to resistance. A more robust method for generating genotype-fitness mappings in an unbiased way will likely be necessary for generalizing these results. Furthermore, the evolutionary model employed here only represents well-mixed *E. coli* growing *in vitro* – we do not model immune system interactions, spatial considerations, or frequency-dependent fitness effects. Finally, we did not explore differential drug penetration into different tissues, which has been found to promote resistance (*48, 58*).

Apart from the specific findings of our work, the fitness seascape model and concept of fitness tradeoffs more generally have larger implications for molecular evolution in varying environments (*26, 59, 60*). The fitness seascape model suggests that selection changes as a function of environmental variables– considering that environmental variables can change over time, selection itself can then change over time. While this is not necessarily a surprising conclusion, this phenomena calls into question the utility of fitness as a scalar, fixed variable. The canonical fitness landscape model in infectious disease posits that higher fitness proteins are those that confer a greater degree of drug resistance, and that evolution will trend towards these higher fitness proteins. In contrast, the fitness seascape model indicates that evolution may not proceed in such a straightforward manner, and selection may be difficult or impossible to predict based on measures of drug resistance alone (*59*). Furthermore, the shape of dose-response curves may change the standing genetic heterogeneity in a population, thus influencing the progression of molecular evolution. For instance, a highly costly mutation that confers a great deal of drug resistance may be exceedingly unlikely to emerge due to selection against the mutation at low drug concentrations. Others have shown that patterns such as global epistasis are also shaped by complex interactions of dose response curves (*61*).

Our work represents the first application of an empirical fitness seascape to predicting the emergence of resistance in the clinic. Our results suggest that explicit modeling of evolutionary trade-offs is important for predicting evolution under time-varying drug concentration, and that this modeling framework may be useful for optimizing drug dosing regimens *in silico* and designing adaptive therapies.

## Materials and Methods

### Model system

Genetically engineered *E. coli* DH5α strains were provided by the Weinreich Lab at Brown University. Details of the genetic engineering are provided by Weinreich et al (*10*). A description of each of the 16 genotypes and their clinical designation, if any, are shown in **Table S1**. We searched NIH BLAST for each β-lactamase sequence to see if it had been previously identified in clinical contexts, noting its clinical designation if applicable.

Cells were cultured from frozen glycerol stock in LB broth. Overnight cultures were grown with 10 μg/mL tetracycline (MP Biomedicals). Cefotaxime (Thermo) was prepared according to manufacturer specifications and stored at -20° C in DI water.

### Growth rate estimation

We generated an optical density to cell count calibration curve using a colony forming unit (CFU) assay. *E. coli* cells were inoculated from frozen glycerol stock in 3 mL LB broth and incubated overnight at 37° C and 220 rpm. A series of 2-fold dilutions with fresh media were made in a 96-well plate and the optical density was measured. Samples corresponding to 1-, 4-, 16-, and 64-fold dilutions were then used to quantify cell count with a CFU assay. Samples were diluted 100,000-fold and 50 μL was deposited on agar plates in triplicate. After overnight culture, individual colonies were counted by eye. We then generated a calibration curve by fitting a linear regression to the log-transformed optical density and cell count data.

To measure the fitness seascape, strains were inoculated into LB broth + 10 *ug*/*mL* tetracycline and grown up overnight in a bacterial shaking incubator set to 37°C and 225 rpm. 16 plates, one plate per genotype, were prepared with a 5-fold cefotaxime dilution scheme and one row free of antibiotic. The plates were organized such that there were 8 experimental replicates per drug concentration. 1 uL of bacteria was inoculated into each well, after which plates were sealed with adhesive film. Plates were then incubated at 37°C with no shaking and transferred by hand to a Tecan Spark microplate reader for OD600 readings every two hours for 20 hours. Plates were shaken at 510 rpm for 10 seconds prior to OD600 readings.

After background subtraction, the time series optical density data were converted to cell count using the calibration curve. We then estimated the growth rate for each condition using an approach adapted from Mira et al (*62*). For each growth curve, the data was log-transformed, the slope of each measurement interval was computed, and the interval containing the maximum rate of change was identified. Intervals surrounding the maximum rate of change were then included for further analysis if their slope was within 0.7x the maximum slope. We then used these identified intervals to compute a linear regression to estimate the growth rate.

To estimate the pharmacodynamic relationship, we fit the following dose-response equation to the estimated growth rates:

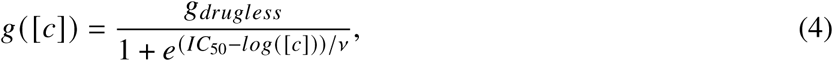

where [*c*] is the drug concentration, *g*_*drugless*_ is the growth rate in the absence of drug, *IC*_50_ is the drug concentration that inhibits the growth rate by 50%, and *v* is the Hill coefficient that determines the steepness of the curve. All data analysis was done in Python using the SciPy package (*63*).

### Genotype- and allele-level analysis

To investigate the relationship between IC_50_ and drug-free growth rate on the scale of individual genotypes, we performed linear regressions on IC_50_ and drug-free growth rate as functions of the total number of drug-resistance mutations. Then, we performed a linear regression on drug free growth rate as a function of IC_50_. To investigate, allele-level effects, we set up a linear regression with each allele and each pairwise interaction term as predictors and either IC_50_ or drug-free growth rate as the outcome. The presence or absence of an allele was coded as a 1 or 0, respectively. Univariate statistical analyses were performed using the SciPy Python package (*63*). Multivariate regressions were performed using the statsmodels Python package (*64*). All data and analysis code can be found at https://doi.org/10.5281/zenodo.14969809.

### Time-kill assay

We developed a novel time-kill assay to estimate drug dose-dependent death rates using alamarBlue cell viability reagent (Thermo).

#### Generating calibration curves

We generated a curve to estimate bacterial cell count from AB fluorescence using a two-step procedure. First, we generated a cell count to optical density calibration curve. *E. coli* cells were inoculated from frozen glycerol stock in 3 mL LB broth and incubated overnight at 37° C and 220 rpm. We made a series of 2-fold dilutions of the culture in fresh media using 90 μL cell culture per well in a clear, polystyrene 96-well plate, with rows B-G serving as technical replicates and columns 2-10 comprising the dilution gradient. Column 11 was used for background estimation by adding fresh medium only. Optical density was measured using a microplate reader, and samples from row B, columns 2, 4, 6, and 8 were subjected to a colony forming unit (CFU) assay to quantify cell count. These samples corresponded to 1-, 4-, 16-, and 64-fold dilutions from the initial culture. We diluted each sample by 100,000-fold and plated 50 μL onto agar plates in triplicate. Colonies were manually counted after overnight incubation at 37° C. We then background-subtracted the optical density data and fit a linear model to the log-transformed optical density and cell count data. We used the slope and y-intercept to estimate cell count from optical density using a linear fit of the log-transformed cell count and OD_600_ values.

To estimate fluorescence, 10 μL AB was added to each well of the same 96-well plate (excluding row B used for the CFU assay) and the plate was incubated for 30 minutes at 37° C with 220 rpm shaking. After incubation, fluorescence was measured with a microplate scanner (excitation filter 540 nm, emission filter 590 nm). After background subtracting, we used the optical density-CFU calibration curve above to estimate cell count as a function of relative fluorescence units (RFU_30_). As before, we estimated the calibration curve with a linear regression on the log-transformed data.

#### Estimation of dose-response curves with time-kill assay

Three 96 well plates were prepared with each plate containing 2 conditions and 3 experimental replicates per condition. Rows B-G were used for experimental replicates while columns 2-11 were used for different timepoints. To optimize the time each condition spent in the dynamic range of the assay, we used different starting concentrations of cells for the two above- and below-MIC conditions. For the above-MIC conditions (conditions where the cell count was not expected to increase), we initialized the experiment with ~2×10^5^ cells/mL, which we obtained by diluting the overnight culture by 4x with fresh media. For the below-MIC conditions, we initialized the experiment with ~2×10^3^ cells/mL by diluting the overnight culture by 100x. All experiments were initialized after 1 hr of pre-incubation after diluting the overnight culture. We prepared a 20,000 μg/mL stock solution of CTX following the manuscturer’s instructions. To ensure that each condition was exposed to drug at roughly the same time, 10 μL of the CTX solution (Thermo) was added to the wells for each condition first, followed by 90 μL of cell culture. The wells on the outer boundary of the plate were left empty, as fluorescence readings were found to be unreliable for these wells.

At time 0, 10 μL AB was added to column 2 of each plate, and plates were incubated for 30 minutes at 37° C with 220 rpm shaking. After 30 minutes, fluorescence was measured with the microplate scanner. 10 μL of AB was added to column three 15 minutes after adding AB to column 2, and plates were incubated for 30 minutes before performing the corresponding fluorescence scan. This process was repeated for columns 4-11, with plates being scanned 30 minutes after adding AB to each column. We used 15-minute time intervals between adding the AB in columns 2-6, 30 minute time intervals for columns 6-8, and 1 hour time intervals for columns 8-11. This approach resulted in sampling time points of roughly 0, 15, 30, 45, 60, 90, 120, 180, 240, and 300 minutes. This sampling scheme allowed us to observe the short- and long-term dynamics of the change in cell count over time. We applied the RFU_30_-cell count calibration curve from above to estimate the cell count at each time point.

#### Net growth rate estimation

Estimating the parameter of interest from time-kill data is a challenge, especially during cell death. While others have used parametric equations constructed from first principles (*65*), we found overfitting to be an impassable challenge. Furthermore, we observed a variable death rate in our high drug concentration conditions. As such, a simple model with fewer parameters may be a more robust approach. As an illustration, we sought to estimate the maximum rate of change in cells using a linear regression on the log-transformed cell count data. To choose a subset of the data that balanced linearity with including as much data as possible, we defined the following objective function:

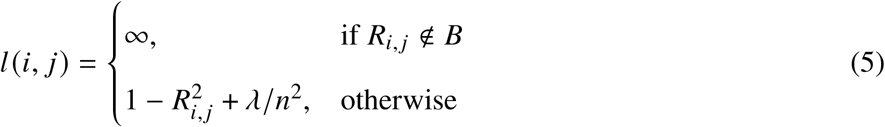

where *i* and *j* are the start and end indices for the subset, *R*_*i*, *j*_ is the correlation coefficient for the subset, 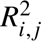 is the coefficient of determination, *n* is the number of points included in the subset, *λ* is a regularization parameter, and *B* is the boundary-defining set. Since 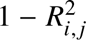 is strictly between 0 and 1, we set *λ* equal to 1. We also defined a boundary set *B* for the estimated slope of the regression based on whether the cell count was increasing or decreasing. For instance, for the drug-free condition, the slope must be greater than 0 since the cell count increased, so *B* = {*x*: *x* > 0}. If *R*_*i*, *j*_ is not in the set *B*, then the loss function is set to infinity. We then minimized *l* (*i*, *j*) for each dataset by an exhaustive search of all subsets larger than 2 elements. For the optimization step, time and cell count for each dataset were normalized between 0 and 1. We then used the subset identified with this process to estimate the maximum rate of change in cell count using a linear regression.

All regressions were performed with the SciPy python package. All code and data required for reproduction is available in the repository: https://doi.org/10.5281/zenodo.14969809.

### Integrated fitness seascape parametrization

We parameterized genotype-specific dose-response curves with the following pharmacodynamic equation:

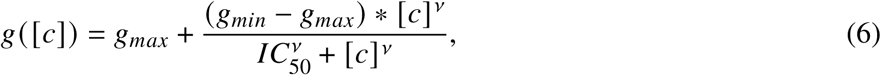

where *g*_*max*_ is the maximum estimated growth rate, *g*_*min*_ is the minimum growth rate (i.e., maximum net loss rate), *IC*_50_ is the half-maximal inhibitory concentration, and *v* is the Hill coefficient. We used time-kill data from genotype 0 to estimate *g*_*min*_ and *v* and applied this to all of the dose-response curves. We used the drug-free growth rate and *IC*_50_ estimated from **Fig 2** to generate genotype-specific dose response curves. These dose-response curves were then used to parametrize FEArS for computational modeling.

### Pharmacokinetic nonadherence modeling

We modeled patient nonadherence by convolving an impulse train representing correctly administered doses with the 2-compartment pharmacokinetic model (**Eq 1**). Formally, this is done by generating a set (*D*) of numbers representing the time of each scheduled drug dose. Then, for each patient, a random subset of drug doses are removed from *D*, with each drug dose having a constant probability *p_forget_* of being removed, resulting in the set of drug doses that the simulated patient correctly self-administered (*D*_*a*_). A Dirac comb is generated using *D*_*a*_ with the period *T* equal to the time between drug doses:

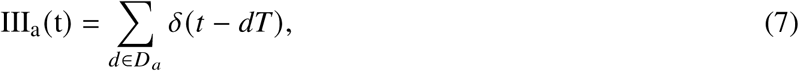

and the resulting impulse train is convolved with the pharmacokinetic model in **Eq 1** to generate the simulated patient’s serum drug concentration:

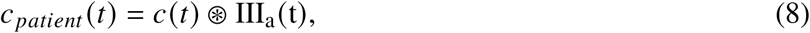

where ⊛ is the convolution operator.

### *In vitro* drug regimen nonadherence model

We developed an *in vitro* model to better understand the impact of early versus late nonadherence. Rows in a 96-well plate B, C, and D were used as experimental conditions, with columns 2-11 being technical replicates. *E. coli* genotypes 0, 1, and 2 were cultured overnight as described previously. 10 μL of genotype 0 culture was added to 90 μL of fresh media in each experimental well for a 1:10 dilution. Cultures of genotypes 1 and 2 were diluted 1:10 in fresh media before adding 1 μL to each well for a 1:10^3^ dilution. Then, wells were dosed with either 10 μg/mL cefotaxime or media control. Every 8 hours, the cells were passaged to a new 96-well plate in a 1:10 dilution and the drug concentration was updated according to the schedule in **Table S2**. Cells were not resuspended in fresh media prior to passaging, so some residual drug was carried over into the subsequent wells. However, this small amount of residual drug represents a difference of at most 1 μg/mL between conditions, and thus does not impact the qualitative result of the experiment. After passaging, 10 μL of AB was added to each well and the cell count was estimated using the method described above. An additional 24 hour AB scan was completed after the 16 hour passage.

## Author contributions

Conceptualization: ESK, DTW, JM, RBC, ED, JGS Methodology: ESK, AES Software: ESK, AES, DTW Validation: ESK, AES Formal analysis: ESK Investigation: ESK, AES Writing–original draft preparation: ESK Writing–review and editing: ESK, AES, DTW, JM, RBC, ED, JGS Visualization: ESK Resources: JGS Supervision: ED, JGS Funding acquisition: JGS

## Data and materials availability

All data and software necessary for replication is freely available at https://doi.org/10.5281/zenodo.14969809.

## Supplementary Materials

### Supplementary Text

#### FEArS: an agent-based modeling platform

We developed FEArS, Fast Evolution on Arbitrary Seascapes, to simulate bacterial populations responding to drug with our fitness seascape and time-kill data. An overview of the core evolutionary algorithm is shown in **Fig. S1**.

#### Mutation supply impacts probability of treatment success

We performed a Monte Carlo simulation using Latin Hypercube Sampling to investigate the impact of maximum serum drug concentration *c*_*max*_, drug half life *t*_1/2_, carrying capacity *K*, and mutation rate *r*_*m*_ on the probability of treatment success in the setting of inpatient *E. coli* bacteremia treated with IV antibiotics. We used a multiple linear regression to analyze the relative importance of each of the parameters on the outcome, shown in **Table S3**. We found that *K* and *r*_*m*_ were both independently and jointly associated with treatment success.

We investigated the impact of the average mutation supply, *Kr*_*m*_, on the probability of treatment success (**Fig S2**). Above ~1 mutant per hour, there was a very low probability of treatment success. We further investigated the genotype-specific cell count for different mutation supply scenarios (**Fig S3**), finding that for higher rates of mutation supply, drug resistant genotypes exist at a low population count in equilibrium with the wild-type.

#### Total administered doses is not a meaningful predictor of treatment success

We sought to determine whether the total number of doses correctly administered may correspond to treatment success (**Fig. S4**). While there are slight differences between the total doses between regimens that resulted in failure and success, the difference in the mean is less than one dose on average, and thus not likely to be clinically significant.

**Table S1:**
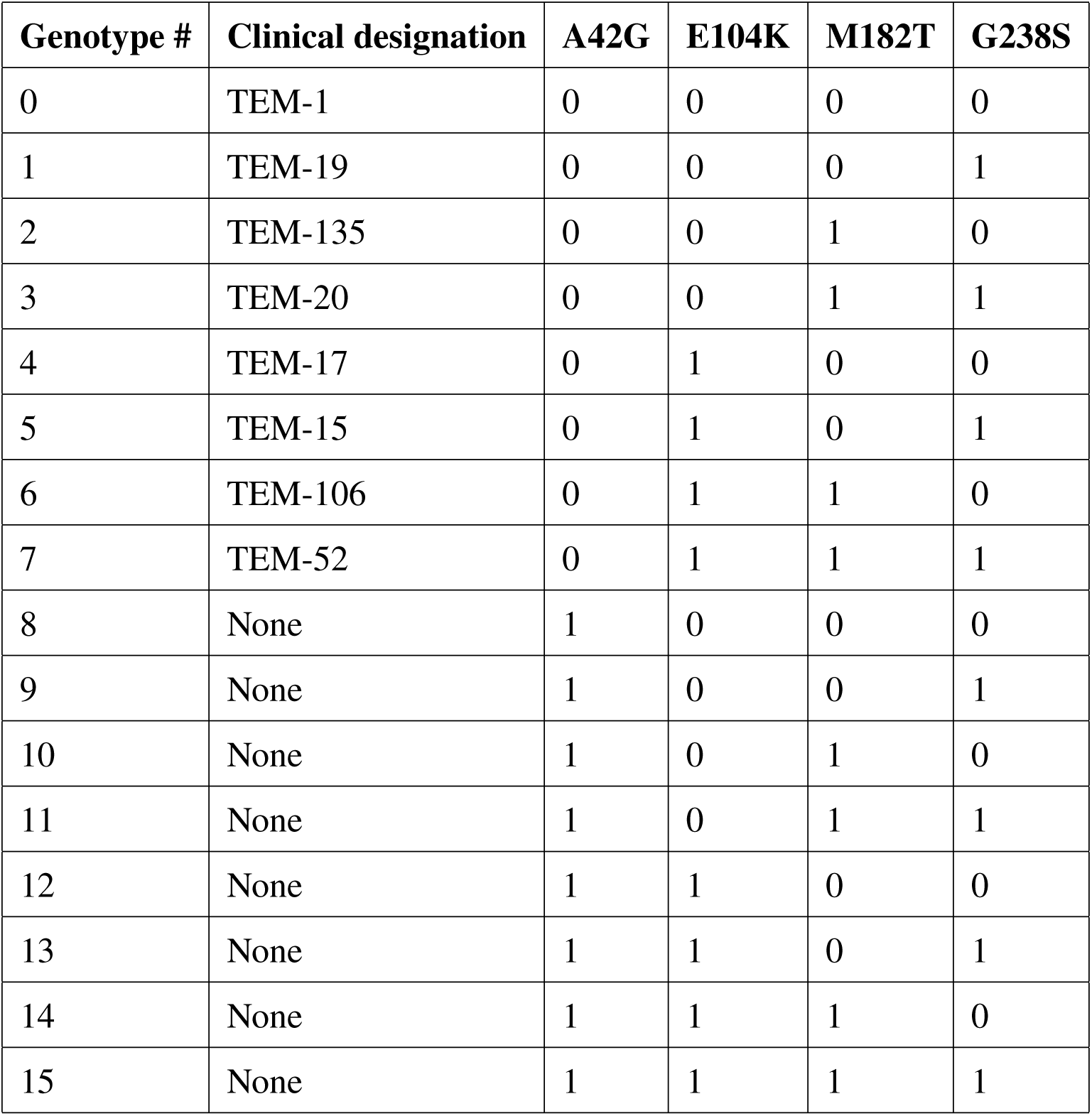
Engineered *E. coli* genotypes and their clinical designations. Columns A42G, E104K, M182T, and G238S each correspond to the four β-lactamase point mutations in the model system. A 1 or 0 in the column for a mutation indicates the presence or absence of that mutation in the genotype.

**Table S2:**
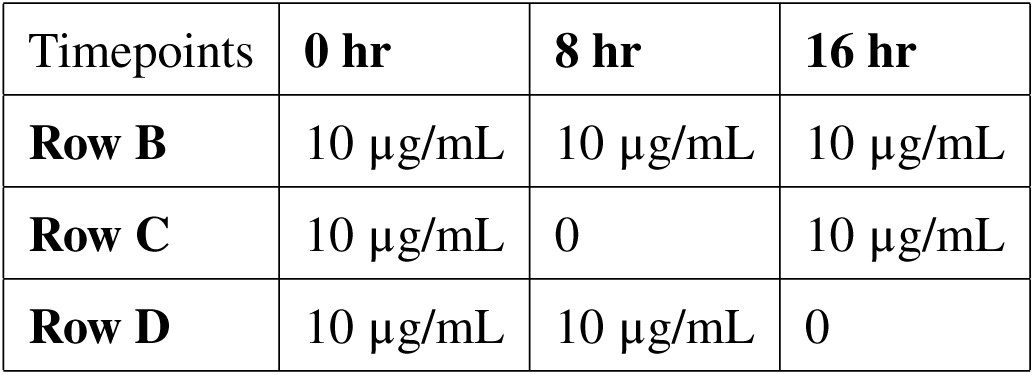
Drug dosing schedule.

**Figure S1:**
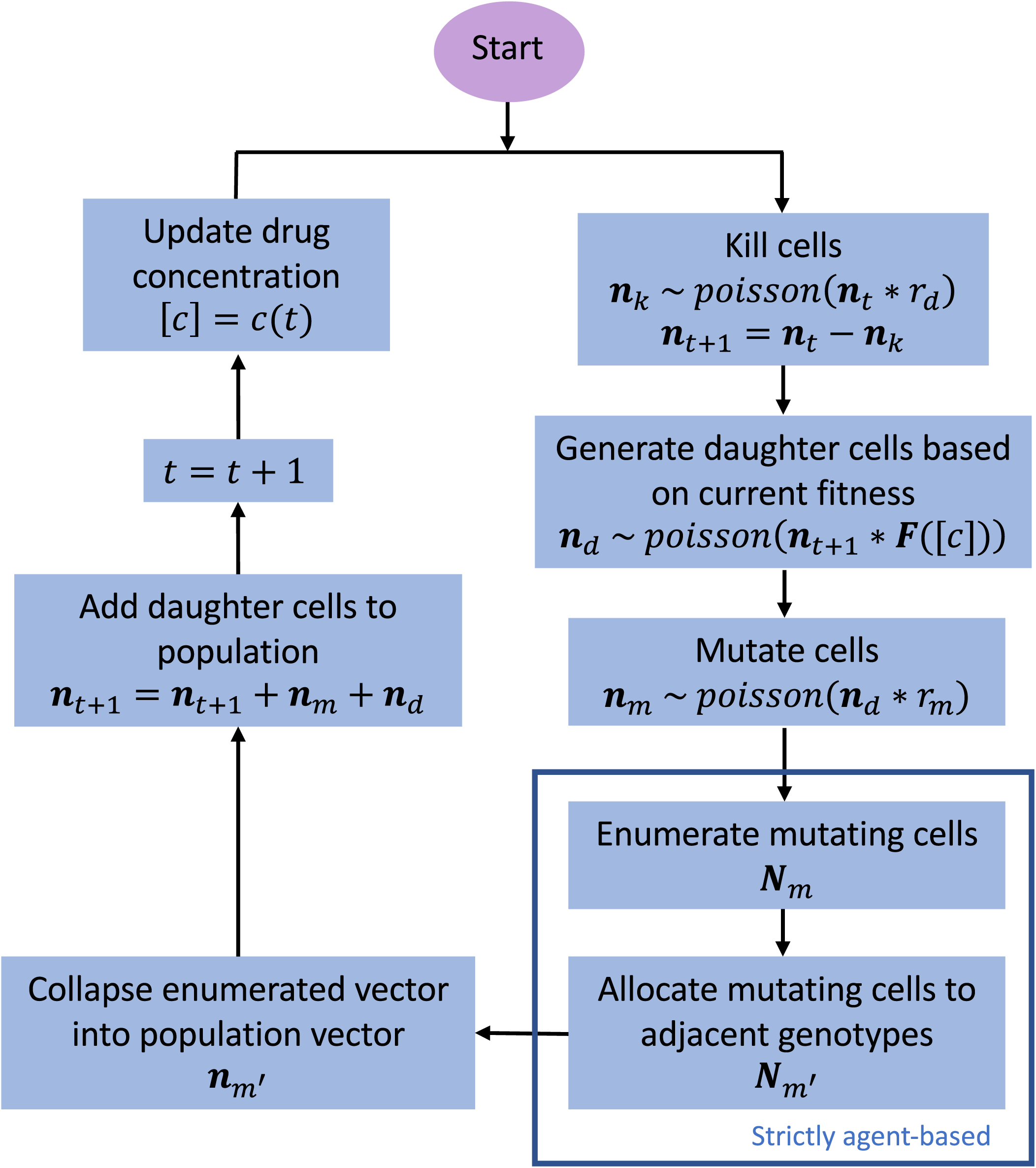
FEArS evolution simulation flowchart.

**Figure S2:**
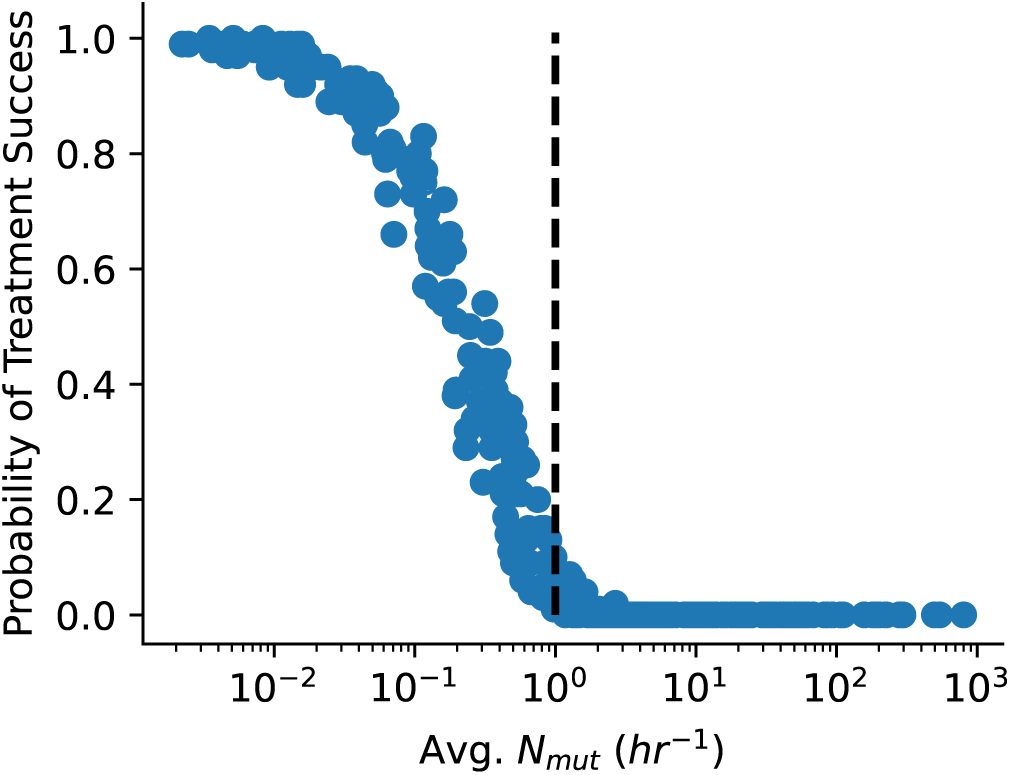
Mutation supply versus probability of treatment success. Mutation supply (Avg *N*_*mut*_) is the product of the mutation rate and the carrying capacity. Vertical line indicates 1 mutant per hour.

**Table S3:** Multiple linear regression analysis and interactions for simulated bacteremia treated with IV antibiotics.

| Variable | Regression Coefficient ($\beta$) | Standard Error |
| --- | --- | --- |
| Intercept | 1.6632 | 0.070 |
| $c_{max}$ | -0.0124 | 0.091 |
| $t_{1/2}$ | -0.3797 | 0.086 |
| $K$ | -1.6488 | 0.087 |
| $r_m$ | -1.6351 | 0.087 |
| $c_{max} \times t_{1/2}$ | 0.0864 | 0.091 |
| $c_{max} \times K$ | -0.0276 | 0.092 |
| $c_{max} \times r_m$ | -0.0680 | 0.095 |
| $t_{1/2} \times K$ | 0.2147 | 0.088 |
| $t_{1/2} \times r_m$ | 0.2399 | 0.090 |
| $K \times r_m$ | 1.4565 | 0.091 |

**Figure S3:**
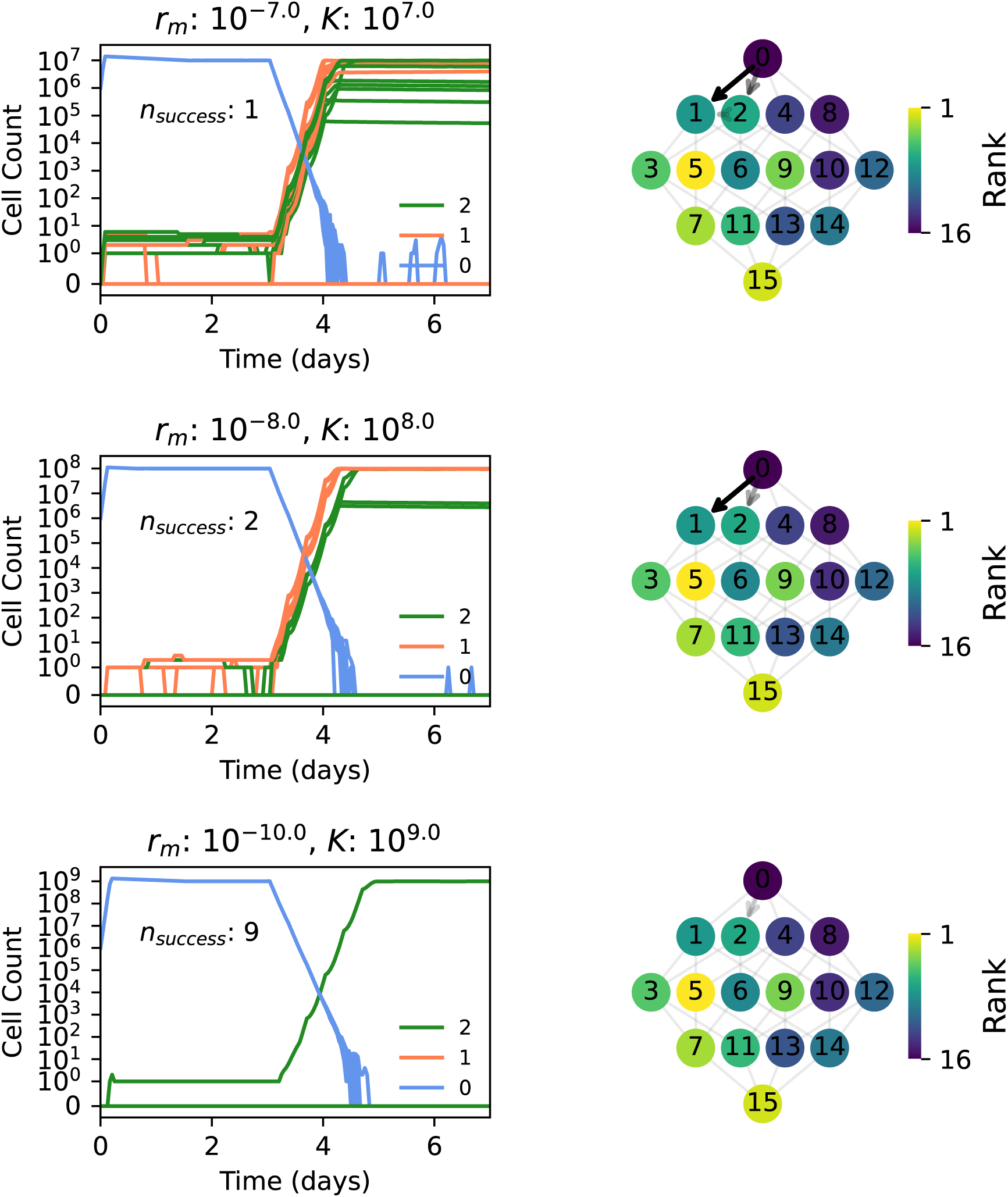
Higher mutation supply increases probability of resistance. Left column: cell count over time for genotypes 0, 1, and 2 for three combinations of *r*_*m*_ and *K*. Top two rows correspond to mutation supply ~1 per cell division; bottom row corresponds to mutation supply ~0.1 per cell division. Right column: evolutionary trajectories corresponding to the data on the left. Arrow opacity corresponds to the number of simulations following a given trajectory, with a maximum of six trajectories for the darkest lines and one trajectory for the lightest lines. *N* = 10 simulations per parameter set; treatment began at *t* = 3 days.

**Figure S4:**
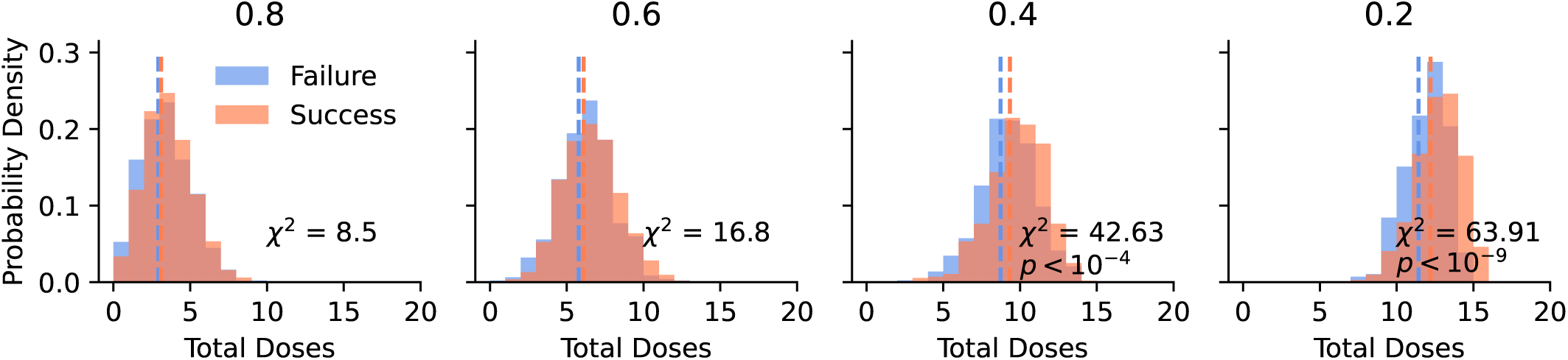
Total number of administered doses is associated with treatment success. Columns correspond to different rates of *p* _*forget*_, labeled at the top. Dotted vertical lines indicate the mean of the distributions. *χ*-squared statistic and p-values are shown inset. *N* = 1000 simulations per *p* _*forget*_.

## References and Notes

1. C. J. L. Murray, et al., Global burden of bacterial antimicrobial resistance in 2019: a systematic analysis. The Lancet 399 (10325), 629–655 (2022), doi:10.1016/S0140-6736(21)02724-0.

2. J. Chmielecki, et al., Optimization of Dosing for EGFR-Mutant Non–Small Cell Lung Cancer with Evolutionary Cancer Modeling. Science Translational Medicine 3 (90) (2011), doi:10.1126/scitranslmed.3002356, https://www.science.org/doi/10.1126/scitranslmed.3002356.

3. S. Chakrabarti, F. Michor, Pharmacokinetics and Drug Interactions Determine Optimum Combination Strategies in Computational Models of Cancer Evolution. Cancer Research 77 (14), 3908–3921 (2017), doi:10.1158/0008-5472.CAN-16-2871.

4. J. Maltas, et al., Drug dependence in cancer is exploitable by optimally constructed treatment holidays. Nature Ecology & Evolution 8 (1), 147–162 (2023), doi:10.1038/s41559-023-02255-x.

5. J. Maltas, K. B. Wood, Pervasive and diverse collateral sensitivity profiles inform optimal strategies to limit antibiotic resistance. PLOS Biology 17 (10), e3000515 (2019), doi:10.1371/journal.pbio.3000515.

6. S. Iram, et al., Controlling the speed and trajectory of evolution with counterdiabatic driving. Nature Physics 17 (1), 135–142 (2021), doi:10.1038/s41567-020-0989-3.

7. A. Fischer, I. Vázquez-Garćıa, V. Mustonen, The value of monitoring to control evolving populations. Proceedings of the National Academy of Sciences 112 (4), 1007–1012 (2015), doi:10.1073/pnas.1409403112.

8. K. E. Poels, et al., Identification of optimal dosing schedules of dacomitinib and osimertinib for a phase I/II trial in advanced EGFR-mutant non-small cell lung cancer. Nature Communications 12 (1), 3697 (2021), doi:10.1038/s41467-021-23912-4.

9. D. T. Weaver, E. S. King, J. Maltas, J. G. Scott, Reinforcement learning informs optimal treatment strategies to limit antibiotic resistance. Proceedings of the National Academy of Sciences 121 (16), e2303165121 (2024), doi:10.1073/pnas.2303165121.

10. D. M. Weinreich, N. F. Delaney, M. A. DePristo, D. L. Hartl, Darwinian Evolution Can Follow Only Very Few Mutational Paths to Fitter Proteins. Science 312 (5770), 111–114 (2006), doi:10.1126/science.1123539.

11. C. P. Goulart, et al., Designing Antibiotic Cycling Strategies by Determining and Understanding Local Adaptive Landscapes. PLoS ONE 8 (2), e56040 (2013), doi:10.1371/journal.pone.0056040.

12. I. Fragata, A. Blanckaert, M. A. Dias Louro, D. A. Liberles, C. Bank, Evolution in the light of fitness landscape theory. Trends in Ecology & Evolution 34 (1), 69–82 (2019), doi:10.1016/j.tree.2018.10.009.

13. R. D. Kouyos, et al., Exploring the Complexity of the HIV-1 Fitness Landscape. PLoS Genetics 8 (3), e1002551 (2012), doi:10.1371/journal.pgen.1002551.

14. J. Maltas, D. M. McNally, K. B. Wood, Evolution in alternating environments with tunable interlandscape correlations. Evolution 75 (1), 10–24 (2021), doi:10.1111/evo.14121.

15. C. J. Watson, et al., The evolutionary dynamics and fitness landscape of clonal hematopoiesis. Science 367 (6485), 1449–1454 (2020), doi:10.1126/science.aay9333.

16. J. H. Gillespie, Molecular Evolution Over the Mutational Landscape. Evolution 38 (5), 1116 (1984), doi:10.2307/2408444.

17. K. M. Brown, et al., Compensatory Mutations Restore Fitness during the Evolution of Dihydrofolate Reductase. Molecular Biology and Evolution 27 (12), 2682–2690 (2010), doi:10.1093/molbev/msq160.

18. M. Lagator, N. Colegrave, P. Neve, Selection history and epistatic interactions impact dynamics of adaptation to novel environmental stresses. Proceedings of the Royal Society B: Biological Sciences 281 (1794), 20141679 (2014), doi:10.1098/rspb.2014.1679.

19. M. S. Costanzo, K. M. Brown, D. L. Hartl, Fitness trade-offs in the evolution of Dihydrofolate reductase and drug resistance in plasmodium falciparum. PLoS One 6 (5) (2011), doi:10.1371/journal.pone.0019636.

20. S. Maisnier-Patin, D. I. Andersson, Adaptation to the deleterious effects of antimicrobial drug resistance mutations by compensatory evolution. Research in Microbiology 155 (5), 360–369 (2004), doi:10.1016/j.resmic.2004.01.019.

21. N. Farrokhian, et al., Measuring competitive exclusion in non–small cell lung cancer. Science Advances 8 (26), eabm7212 (2022), doi:10.1126/sciadv.abm7212.

22. M. Lässig, V. Mustonen, A. M. Walczak, Predicting evolution. Nature Ecology & Evolution 1 (3), 0077 (2017), doi:10.1038/s41559-017-0077.

23. D. J. Merrell, The Adaptive Seascape: The Mechanism of Evolution (University of Minnesota Press), minne ed. edition ed. (1994).

24. C. B. Ogbunugafor, C. S. Wylie, I. Diakite, D. M. Weinreich, D. L. Hartl, Adaptive Landscape by Environment Interactions Dictate Evolutionary Dynamics in Models of Drug Resistance. PLOS Computational Biology 12 (1), e1004710 (2016), doi:10.1371/journal.pcbi.1004710.

25. S. G. Das, S. O. Direito, B. Waclaw, R. J. Allen, J. Krug, Predictable properties of fitness landscapes induced by adaptational tradeoffs. eLife 9, e55155 (2020), doi:10.7554/eLife.55155.

26. V. Mustonen, M. Lässig, From fitness landscapes to seascapes: non-equilibrium dynamics of selection and adaptation. Trends in Genetics 25 (3), 111–119 (2009), doi:10.1016/j.tig.2009.01.002.

27. A. Agarwala, D. S. Fisher, Adaptive walks on high-dimensional fitness landscapes and seascapes with distance-dependent statistics. Theoretical Population Biology 130, 13–49 (2019), doi:10.1016/j.tpb.2019.09.011.

28. E. S. King, D. S. Tadele, B. Pierce, M. Hinczewski, J. G. Scott, Diverse mutant selection windows shape spatial heterogeneity in evolving populations. PLOS Computational Biology 20 (2), e1011878 (2024), doi:10.1371/journal.pcbi.1011878.

29. D. Pittet, N. Li, R. P. Wenzel, Association of secondary and polymicrobial nosocomial bloodstream infections with higher mortality. European Journal of Clinical Microbiology & Infectious Diseases 12 (11), 813–819 (1993), doi:10.1007/BF02000400.

30. A. Gikas, et al., Gram-negative bacteremia in non-neutropenic patients: A 3-year review. Infection 26 (3), 155–159 (1998), doi:10.1007/BF02771841.

31. J. Fitzpatrick, et al., Gram-negative bacteraemia; a multi-centre prospective evaluation of empiric antibiotic therapy and outcome in English acute hospitals. Clinical Microbiology and Infection 22 (3), 244–251 (2016), doi:10.1016/j.cmi.2015.10.034.

32. M. K. Singh, B. N. Dominy, The Evolution of Cefotaximase Activity in the TEM β-Lactamase. Journal of Molecular Biology 415 (1), 205–220 (2012), 10.1016/j.jmb.2011.10.041, https://www.sciencedirect.com/science/article/pii/S0022283611011892.

33. K. B. Patel, D. P. Nicolau, C. H. Nightingale, R. Quintiliani, Pharmacokinetics of cefotaxime in healthy volunteers and patients. Diagnostic Microbiology and Infectious Disease 22 (1–2), 49–55 (1995), doi: 10.1016/0732-8893(95)00072-I.

34. I. Matic, et al., Highly Variable Mutation Rates in Commensal and Pathogenic Escherichia coli. Science 277 (5333), 1833–1834 (1997), doi:10.1126/science.277.5333.1833.

35. P. D. Sniegowski, P. J. Gerrish, R. E. Lenski, Evolution of high mutation rates in experimental populations of E. coli. Nature 387 (6634), 703–705 (1997), doi:10.1038/42701.

36. M. D. McKay, R. J. Beckman, W. J. Conover, A Comparison of Three Methods for Selecting Values of Input Variables in the Analysis of Output from a Computer Code. Technometrics 21 (2), 239 (1979), doi:10.2307/1268522.

37. J. E. LeClerc, B. Li, W. L. Payne, T. A. Cebula, High Mutation Frequencies Among Escherichia coli and Salmonella Pathogens. Science 274 (5290), 1208–1211 (1996), doi:10.1126/science.274.5290.1208.

38. P. Komp Lindgren, r. Karlsson, D. Hughes, Mutation Rate and Evolution of Fluoroquinolone Resistance in Escherichia coli Isolates from Patients with Urinary Tract Infections. Antimicrobial Agents and Chemotherapy 47 (10), 3222–3232 (2003), doi:10.1128/AAC.47.10.3222-3232.2003.

39. M. Fernandes, et al., Non-adherence to antibiotic therapy in patients visiting community pharmacies. International Journal of Clinical Pharmacy 36 (1), 86–91 (2014), doi:10.1007/s11096-013-9850-4.

40. F. D’Ambrosio, et al., Adherence to Antibiotic Prescription of Dental Patients: The Other Side of the Antimicrobial Resistance. Healthcare 10 (9), 1636 (2022), doi:10.3390/healthcare10091636.

41. B. A. Almomani, B. M. Hijazi, O. Awwad, R. A. Khasawneh, Prevalence and predictors of non-adherence to short-term antibiotics: A population-based survey. PLOS ONE 17 (5), e0268285 (2022), doi:10.1371/journal.pone.0268285.

42. A. H. Melnyk, A. Wong, R. Kassen, The fitness costs of antibiotic resistance mutations. Evolutionary Applications 8 (3), 273–283 (2015), doi:10.1111/eva.12196.

43. V. Sideraki, W. Huang, T. Palzkill, H. F. Gilbert, A secondary drug resistance mutation of TEM-1 β-lactamase that suppresses misfolding and aggregation. Proceedings of the National Academy of Sciences 98 (1), 283–288 (2001), doi:10.1073/pnas.98.1.283.

44. T. Palzkill, Structural and Mechanistic Basis for Extended-Spectrum Drug-Resistance Mutations in Altering the Specificity of TEM, CTX-M, and KPC β-lactamases. Frontiers in Molecular Biosciences 5, 16 (2018), doi:10.3389/fmolb.2018.00016.

45. A. Nande, A. L. Hill, The risk of drug resistance during long-acting antimicrobial therapy. Proceedings of the Royal Society B: Biological Sciences 289 (1986), 20221444 (2022), doi:10.1098/rspb.2022.1444.

46. M. Lipsitch, B. R. Levin, The population dynamics of antimicrobial chemotherapy. Antimicrobial Agents and Chemotherapy 41 (2), 363–373 (1997), doi:10.1128/AAC.41.2.363.

47. D. I. S. Rosenbloom, A. L. Hill, S. A. Rabi, R. F. Siliciano, M. A. Nowak, Antiretroviral dynamics determines HIV evolution and predicts therapy outcome. Nature Medicine 18 (9), 1378–1385 (2012), doi:10.1038/nm.2892.

48. A. F. Feder, K. N. Harper, C. J. Brumme, P. S. Pennings, Understanding patterns of HIV multi-drug resistance through models of temporal and spatial drug heterogeneity. eLife 10, e69032 (2021), doi: 10.7554/eLife.69032.

49. R. D. Kouyos, P. Abel zur Wiesch, S. Bonhoeffer, Informed Switching Strongly Decreases the Prevalence of Antibiotic Resistance in Hospital Wards. PLoS Computational Biology 7 (3), e1001094 (2011), doi: 10.1371/journal.pcbi.1001094.

50. P. M. Mira, et al., Rational Design of Antibiotic Treatment Plans: A Treatment Strategy for Managing Evolution and Reversing Resistance. PLOS ONE 10 (5), e0122283 (2015), doi:10.1371/journal.pone.0122283.

51. D. Nichol, et al., Steering Evolution with Sequential Therapy to Prevent the Emergence of Bacterial Antibiotic Resistance. PLOS Computational Biology 11 (9), e1004493 (2015), doi:10.1371/journal.pcbi.1004493.

52. D. Nichol, et al., Antibiotic collateral sensitivity is contingent on the repeatability of evolution. Nature Communications 10 (1), 334 (2019), doi:10.1038/s41467-018-08098-6.

53. S. Kim, T. D. Lieberman, R. Kishony, Alternating antibiotic treatments constrain evolutionary paths to multidrug resistance. Proceedings of the National Academy of Sciences 111 (40), 14494–14499 (2014), doi:10.1073/pnas.1409800111.

54. P. Abel zur Wiesch, R. Kouyos, S. Abel, W. Viechtbauer, S. Bonhoeffer, Cycling Empirical Antibiotic Therapy in Hospitals: Meta-Analysis and Models. PLoS Pathogens 10 (6), e1004225 (2014), doi:10.1371/journal.ppat.1004225.

55. S. Iram, et al., Controlling the speed and trajectory of evolution with counterdiabatic driving. Nature Physics 17 (1), 135–142 (2021).

56. A. J. Claxton, J. Cramer, C. Pierce, A systematic review of the associations between dose regimens and medication compliance. Clinical Therapeutics 23 (8), 1296–1310 (2001), doi:10.1016/S0149-2918(01)80109-0.

57. P. Kardas, Patient compliance with antibiotic treatment for respiratory tract infections. Journal of Antimicrobial Chemotherapy 49 (6), 897–903 (2002), doi:10.1093/jac/dkf046.

58. S. Moreno-Gamez, et al., Imperfect drug penetration leads to spatial monotherapy and rapid evolution of multidrug resistance. Proceedings of the National Academy of Sciences 112 (22), E2874–E2883 (2015), doi:10.1073/pnas.1424184112.

59. A.-H. Ghenu, A. Amado, I. Gordo, C. Bank, Epistasis decreases with increasing antibiotic pressure but not temperature. Philosophical Transactions of the Royal Society B: Biological Sciences 378 (1877), 20220058 (2023), doi:10.1098/rstb.2022.0058.

60. H. A. Lindsey, J. Gallie, S. Taylor, B. Kerr, Evolutionary rescue from extinction is contingent on a lower rate of environmental change. Nature 494 (7438), 463–467 (2013), doi:10.1038/nature11879.

61. J. Diaz-Colunga, A. Sanchez, C. B. Ogbunugafor, Environmental modulation of global epistasis in a drug resistance fitness landscape. Nature Communications 14 (1), 8055 (2023), doi:10.1038/s41467-023-43806-x.

62. P. Mira, M. Barlow, J. C. Meza, B. G. Hall, Statistical Package for Growth Rates Made Easy. Molecular Biology and Evolution 34 (12), 3303–3309 (2017), doi:10.1093/molbev/msx255.

63. P. Virtanen, et al., SciPy 1.0: Fundamental Algorithms for Scientific Computing in Python. Nature Methods 17, 261–272 (2020), doi:10.1038/s41592-019-0686-2.

64. S. Seabold, J. Perktold, statsmodels: Econometric and statistical modeling with python, in 9th Python in Science Conference (2010).

65. R. C. Li, Simultaneous pharmacodynamic analysis of the lag and bactericidal phases exhibited by betalactams against Escherichia coli. Antimicrobial Agents and Chemotherapy 40 (10), 2306–2310 (1996), doi:10.1128/AAC.40.10.2306.

